# The luminal and cytosolic domains of CLIMP-63 cooperate to set ER sheet width

**DOI:** 10.64898/2026.06.30.735628

**Authors:** Arthur Samurkas, Laurence Abrami, Béatrice Kunz, Sylvia Ho, Matteo Dal Peraro, Franciso S. Mesquita, F. Gisou van der Goot

## Abstract

The endoplasmic reticulum (ER) synthesized and folds secreted and transmembrane protein in regions composed of flattened sheets with a defined luminal width of ∼50 nm. CLIMP-63 has been proposed to be the principal determinant of this spacing, but the molecular mechanism by which it controls luminal width remains unclear. The luminal domain of CLIMP-63 has been proposed to act as a fixed-length spacer through antiparallel coiled-coil dimerization between molecules on opposing membranes. Here we revise this model. Using systematic cysteine trapping across the luminal domain, we demonstrate that CLIMP-63 assembles into parallel trimers. An AlphaFold-guided screen of CLIMP-63 orthologs combined with cryo-electron microscopy reveals that the luminal domain forms an elongated trimeric rod with intrinsic conformational flexibility at hinge regions. This flexibility is functionally required: replacing the human luminal domain with a more rigid ortholog causes complete ER vacuolation. Higher-order assembly of CLIMP-63 trimers beyond the trimer depends on the cytosolic tail rather than on luminal trans interactions, and the tail’s role in higher order assembly is separable from its other architectural functions. Together, these findings show that luminal trimeric extension and cytosolic tail-mediated clustering cooperate to determine ER sheet width.

## INTRODUCTION

The endoplasmic reticulum (ER) is the largest organelle in most cells, forming a continuous membrane network that extends from the nuclear envelope throughout the cytoplasm. Two major structural domains can be distinguished: a peripheral network of tubules interconnected by three-way junctions, and a perinuclear region of flattened cisternae, termed ER sheets. Both are maintained by dedicated membrane-shaping proteins. Tubule curvature is stabilized by reticulon and DP1/Yop1 family proteins, which oligomerize at high curvature to drive and maintain tubule geometry [1–6]. Sheet morphology is less well understood, but requires luminal spacer proteins and is influenced by the cytoskeleton, other organelles, and by the organization of ribosomes on the outer membrane surface [4,7–14].

ER sheets are the primary site of secretory and membrane protein synthesis. Their outer surface is densely coated with ribosomes organized into polysomes and the lumen accommodates folding chaperones, and quality control factors that process newly synthesized proteins co-translationally. The luminal width, i.e. the distance between the two opposing membrane leaflets, is remarkably constant across cell types, approximately 50 nm in most mammalian cells [4,14–16], though the functional consequences of luminal width regulation are not yet fully understood.

CLIMP63 (cytoskeleton-linking membrane protein of 63 kDa; also known as CKAP4) is a key regulator of ER sheet morphology and luminal width. It is a type II transmembrane protein with a short N-terminal cytoplasmic tail and a large C-terminal luminal domain [4,15,17,18]. Overexpression of CLIMP63 promotes ER sheet formation, while its depletion shifts the balance toward a more tubular morphology [4]. The cytoplasmic tail contributes to ER organization through microtubule binding, which links the ER to the cytoskeleton [11,19–21], and is regulated by phosphorylation and S-acylation [17,21,22].

The luminal domain of CLIMP63 was proposed to act as a lumenal spacer through antiparallel coiled-coil dimerization: molecules anchored in opposing ER membranes would dimerize across the lumen, holding the two membranes at a fixed distance set by the length of the coiled-coil [4,14,23]. However, the antiparallel dimer model raises mechanistic difficulties. Antiparallel arrangements are geometrically disfavored in homo-oligomeric coiled-coils, and direct biochemical evidence for stable trans interactions between CLIMP-63 molecules on opposing membranes has remained elusive.

We recently showed that the CLIMP63 luminal domain, when provided with a signal sequence to direct it to the ER (CLIMP63^LD^), is secreted by cells as a trimer [17]. These findings cannot be reconciled with the antiparallel dimer model and suggest instead that the luminal domain forms a parallel trimeric coiled-coil, consistent with the geometry of canonical homo-trimeric coiled-coil assemblies.

Here we characterize the CLIMP63 luminal domain assembly at the molecular level. Using systematic cysteine-trapping across the length of the luminal domain, we demonstrate that CLIMP-63 forms parallel trimers, and that the trimer is the elementary unit throughout the protein’s cellular itinerary. Using AlphaFold-guided screening of CLIMP-63 orthologs and single-particle cryo-electron microscopy, we establish that the luminal domain forms an elongated trimeric rod with regions of intrinsic conformational flexibility. Through chimeric protein analysis, we show that this flexibility is required for ER architecture: its replacement with a more rigid ortholog domain causes complete ER vacuolation. Finally, using pharmacological perturbations, truncation analysis, and Blue Native PAGE, we demonstrate that trimers associate into higher-order assemblies through their cytoplasmic tails, independently of microtubule binding, S-acylation, and nucleic acid interactions, and that tail-mediated oligomerization is functionally separable from the tail’s other roles in ER architecture. Together, these findings provide a revised structural and mechanistic foundation for understanding how CLIMP-63 shapes the ER lumen.

## RESULTS

### CLIMP63 forms parallel trimers throughout its luminal domain

CLIMP63 is a type II membrane protein with its large C-terminal domain residing in the ER lumen. The luminal domain has been proposed to regulate ER sheet spacing through antiparallel coiled-coil-mediated dimerization between molecules on opposing membranes. We have however recently shown that the isolated CLIMP63 luminal domain, equipped with a signal sequence to direct it to the ER lumen CLIMP63^LD^), is secreted as a trimer [17]. In that study, we also showed by Blue Native PAGE that full-length CLIMP63 assembles into higher-order complexes beyond the trimer, and that data-driven mathematical modelling of the CLIMP63 life cycle required the assembly of elementary trimeric units into larger complexes to accurately capture the experimental observations [17]. Importantly, a trimeric assembly is consistent with a parallel rather than antiparallel coiled-coil organization: standard homo-trimeric coiled-coils are parallel structures, and an antiparallel arrangement between three chains is geometrically disfavored.

To further characterize trimerization, we used AlphaFold to generate a model of a trimer of the transmembrane and luminal domains (Figure 1A). AlphaFold predicts a coiled-coil assembly interrupted by two flexible regions, between residues 383 and 390 and between residues 522 and 526. Due to the extent of the second flexible region, AlphaFold proposes that the distal coiled-coil segment folds back onto the proximal segment.

**Figure 1:**
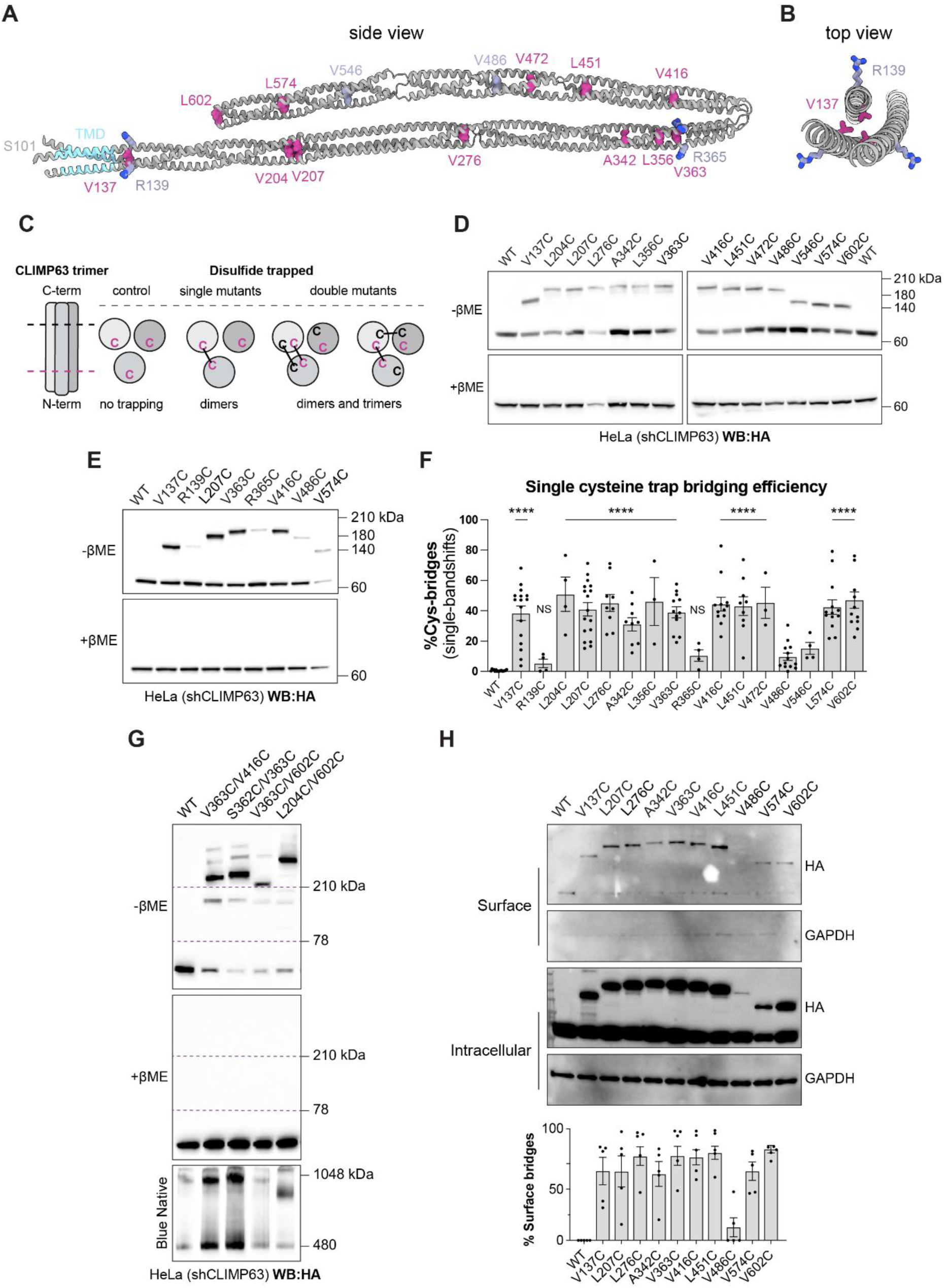
CLIMP63 forms parallel trimers throughout its luminal domain. **A.** Protein structure of trimeric human CLIMP63 (residues 101-602) predicted by AlphaFold3. Amino acid mutated to cysteines are represented. Cyan highlights the transmembrane domain (residues 109-131). The positions oriented towards the interior of the coiled-coil for which cysteine mutagenesis led to efficient disulfide bond formation are shown in pink. Purple color indicated positions that have low cysteine trapping efficiency, with the arginines oriented towards the outside of the colied-coil. **B.** Top view of **(A)**. **C.** Schematic representation of cysteine trapping experiments. Residues mutated into a cysteine that are close enough will form disulfide bridges. Single mutants will generate dimers, whereas double cysteine mutants can either make dimers, trimers, or potentially higher-order interactions. The illustration was created with BioRender.com. **D.** shCLIMP63 HeLa cells overexpressing HA-CLIMP63 WT or single cysteine mutants. Western bloth shows non-reducing (+βME) and reducing (-βME) gels probed for anti-HA. **E.** same as in **(d)** including control mutants R139C and R365C. **F.** Quantification of CLIMP63 cysteine bridge efficiency as in **(d)** and **(E).** Error bars represent mean ± standard deviation (n=3-17). ****: p<0.0001. **G.** shCLIMP63 HeLa cells overexpressing HA-CLIMP63 WT or double cysteine mutants. Western bloth shows non-reducing (+βME) and reducing (-βME) gels probed for anti-HA. Bottom panel shows Western blot analysis of CLIMP63 WT and double cysteine mutants migrated on Blue Native PAGE gel. **H.** Western blot of surface biotinylated proteins and flowthrough (intracellular) from shCLIMP63 HeLa cells transfected with HA-CLIMP63 WT and single cysteine mutants. Surface and intracellular fraction were immunoblotted for HA, with GAPDH used as a cytosolic control. Surface CLIMP63 was quantified by densitometry. Error bars represent mean ± standard deviation (n=5). ****: p<0.0001

To experimentally probe the orientation of CLIMP63 molecules within the trimer, we designed luminal cysteine-trapping mutants by substituting residues predicted to face the interior of the coiled-coil with cysteine (Figure 1B). In a parallel trimer, residues at the same sequence position in adjacent chains are in close proximity and could potentially form disulfide bonds; in an antiparallel arrangement, equivalent positions would be far apart and crosslinking would not be expected. As controls, we substituted residues predicted to face outward. Mutants were expressed in shCLIMP63 HeLa cells and disulfide bond formation was assessed by non-reducing SDS-PAGE. All single cysteine substitutions at interior-facing positions, albeit with reduced efficiency for V486C and V546C, generated a higher molecular weight band (Figure 1C,D). Since each molecule carries only one cysteine, a disulfide can only form between two molecules, confirming that these bands are dimers. The apparent mobility of the dimer bands varied with the position of the cysteine along the domain: mutants in the central region migrated more slowly than those near the membrane-proximal or distal ends. This can be explained geometrically: a crosslink in the middle of two parallel chains creates an extended, X-shaped dimer that could migrate more slowly than a crosslink near either end, which produces an end-stacked arrangement. Control substitutions at exterior-facing positions showed markedly lower crosslinking efficiency, comparable to V486C confirming the specificity of the approach (Figure 1E,F). Thus the inner-facing positions that support efficient crosslinking are consistent with a parallel, in-register arrangement.

We next introduced double cysteine substitutions. In all cases, double mutants generated a predominant high molecular weight band larger than any dimer observed with single mutants, consistent with a crosslinked trimer (Figure 1G). Remarkably few species larger than trimers were detected, consistent with the trimer being the elementary CLIMP63 assembly unit [17]. This was confirmed using CLIMP63^LD^: double and triple cysteine CLIMP63^LD^ mutants migrated as a single high molecular weight band on non-reducing SDS-PAGE (Fig. S1A), demonstrating that the trimer is crosslinkable throughout the luminal domain.

Blue Native PAGE analysis of crosslinked CLIMP63 mutants showed a migration pattern similar to that of wild-type CLIMP63, with two prominent bands at apparent molecular weights of ∼480 and ∼1,048 kDa (Figure 1G), confirming that disulfide trapping does not alter higher-order assembly. The ∼1,048 kDa species was absent when analyzing CLIMP63^LD^, whether wild-type or crosslinked (Fig. S1A), indicating that this species requires elements outside the luminal domain.

Finally, surface biotinylation followed by streptavidin pulldown and non-reducing western blotting showed that disulfide-crosslinked species reach the plasma membrane (Figure 1H). Since the largest species detected by cysteine trapping are trimers, the presence of crosslinked trimers at the cell surface indicates that the trimer constitutes the elementary unit of CLIMP63 throughout its entire cellular itinerary, from the ER to the plasma membrane.

**Figure S1:**
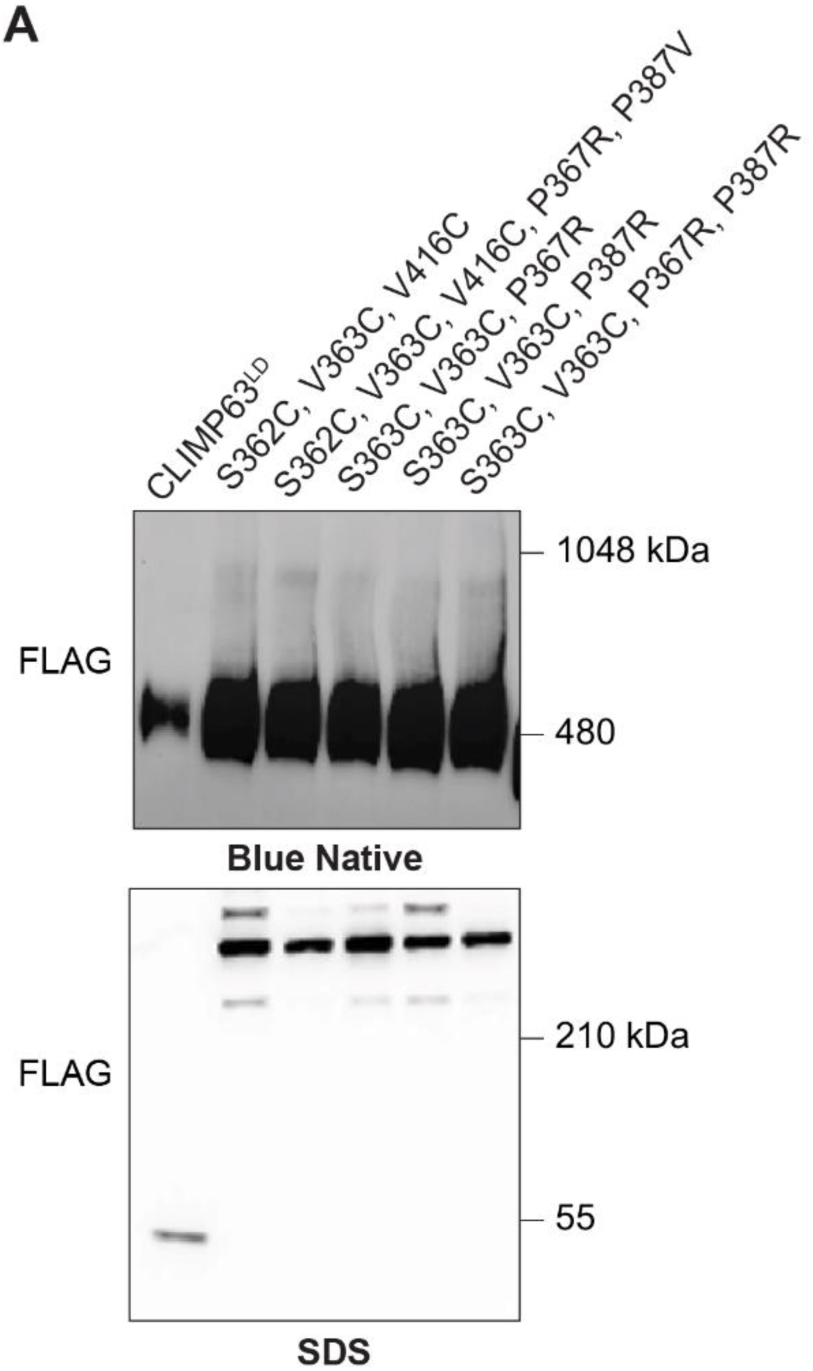
CLIMP63 forms parallel trimers throughout its luminal domain. **A.** Blue Native gel analysis of purified His6x-FLAG-tagged CLIMP63LD and mutants harvested from the cultute medium of suspension-adapted HEK293E cell, immunoblotted for anti-FLAG. **Bottom.** Western blot analysis of purified His6x-FLAG-tagged CLIMP63LD and mutants harvested from the cultute medium of suspension-adapted HEK293E cells, immunoblotted for anti-FLAG.

### CLIMP63 LD has a trimeric rod-shaped alpha-helical structure

The AlphaFold model of trimeric CLIMP63 predicts two regions of conformational flexibility within the coiled-coil. Such flexibility introduces conformational heterogeneity that hinders structural determination by single-particle cryo-electron microscopy, since averaging requires particles to adopt a limited number of discrete conformations. To identify a more tractable model system, we performed an AlphaFold screen of CLIMP63 from 326 species to identify orthologs predicted to have fewer flexible regions. Structures were ranked according to overall helicity, model confidence (pLDDT), predicted aligned error (PAE), and the number of residues in loop regions (Figure 2A). Fish CLIMP63 orthologs generally ranked favorably by these criteria; milkfish CLIMP63 in particular showed fewer predicted flexible regions and a model confidence above 70% throughout most of the domain (Figure 2B).

**Figure 2:**
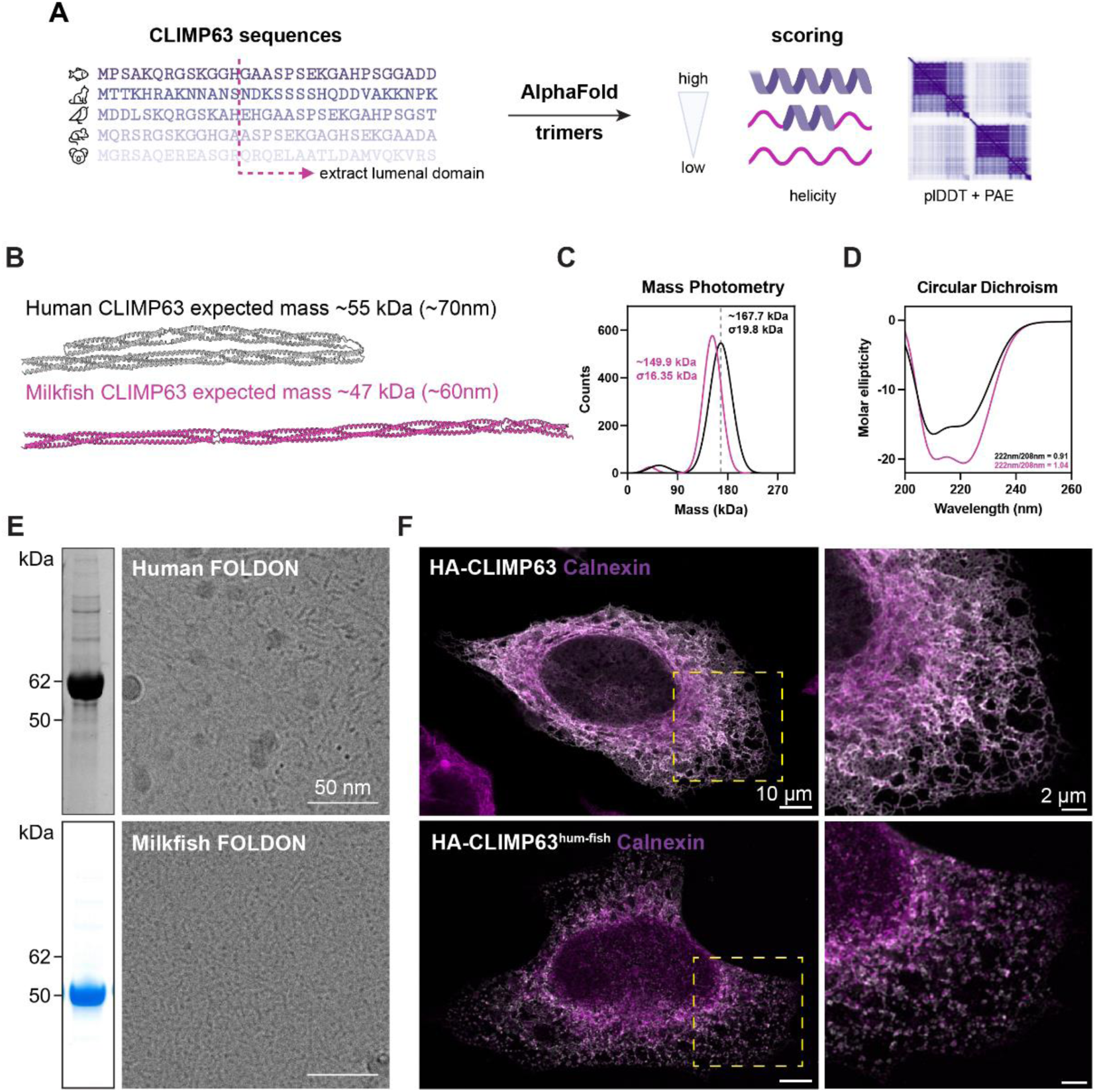
CLIMP63 LD has a trimeric rod-shaped alpha-helical structure. **A.** Schematic illustrating the AlphaFold3 screening. CLIMP63 sequences were obtained from UNIPROT. The transmembrane domain of each sequence was determined using DeepTMHMM [35]. Protein structures were predicted by AlphaFold3 as trimers were ranked according to overall helicity, model confidence (pLDDT), predicted aligned error (PAE), and the number of residues in loop regions. **B.** Protein structure of human and milkfish predicted by AlphaFold3 as trimers. **C.** Mass photometry of purified human and milkfish His6x-FLAG-tagged CLIMP63LD. **D.** Circular Dichroism (CD) spectrum of purified human and milkfish His6x-FLAG-tagged CLIMP63LD. **E.** Coomassie Blue staining of human and milkfish His6x-FLAG-tagged FOLDON-CLIMP63LD. Representative cryo-EM micrographs of both constructs. Scale bar: 50 nm **f** Immunofluorescence of shCLIMP63 HeLa cells expressing human HA-CLIMP63, and CLIMP63hum-fish, labelled for HA (Grey) and ER marker Calnexin (Magenta) Scale bar: 10 μm and 2 μm.

To confirm that milkfish CLIMP63 also forms trimers, we expressed and purified the FLAG-tagged luminal domain (milkfish CLIMP63^LD^). Mass photometry showed that milkfish CLIMP63LD is trimeric (Figure 2C), and circular dichroism confirmed an exclusively alpha-helical secondary structure (Figure 2D), consistent with an extended coiled-coil.

To improve particle behavior for structural studies, we fused human and milkfish CLIMP63^LD^ to the bacteriophage T4 fibritin foldon domain [24,25], which enforces trimerization and provides a rigid cap that promotes favorable particle orientation in vitrified ice. Cryo-EM analysis revealed rod-shaped particles for both constructs (Figure 2E). The apparent length of human foldon-CLIMP63LD particles was shorter than predicted by the AlphaFold model, consistent with partial collapse of the coiled-coil at one or both hinge regions. In contrast, milkfish foldon-CLIMP63LD particles showed dimensions consistent with the AlphaFold prediction and adopted more favorable orientations in the ice layer. Together, these data establish that the CLIMP63 luminal domain forms an elongated trimeric alpha-helical structure, fully extended in milkfish.

### Rigidifying the CLIMP63 luminal domain disrupts ER architecture

To test whether the flexibility of the human CLIMP63 luminal domain is functionally important, we generated a chimeric protein harboring the N-terminal cytoplasmic tail and transmembrane domain of human CLIMP63 fused to the luminal domain of milkfish CLIMP63 (CLIMP63^hum-fish^). Human CLIMP63 or CLIMP63^hum-fish^ were expressed in shCLIMP63 HeLa cells, in which endogenous CLIMP63 is stably depleted. Expression of human CLIMP63 rescued normal ER architecture, whereas expression of CLIMP63^hum-fish^ induced complete vacuolation of the ER (Figure 2F). The only characterized luminal binding partner of CLIMP63 is calumenin, which acts as a chaperone for CLIMP63 rather than as a structural co-organizer of ER sheets [14]; no luminal interactions between CLIMP63 and other ER shaping proteins have been documented. The vacuolation phenotype is therefore unlikely to reflect incompatibility of the milkfish domain with human luminal binding partners, and instead suggests that the conformational flexibility conferred by the hinge regions of the human luminal domain is itself required for the ER-shaping function of CLIMP63.

### CLIMP63 trimers associate into higher-order structures through their cytosolic tails

Blue Native PAGE of CLIMP63 resolves two prominent bands at apparent molecular weights of ∼480 and ∼1,048 kDa [17], Figure 1G and 3B). Since the true molecular weight of a CLIMP63 trimer is approximately 190 kDa, the apparent sizes reflect contributions from bound detergent and lipid as well as the elongated shape of the trimeric molecule. On the same basis, the ∼1,048 kDa band, which migrates at approximately twice the apparent mass of the ∼480 kDa species, likely corresponds to a higher-order assembly, tentatively a dimer of trimers.

**Figure 3.**
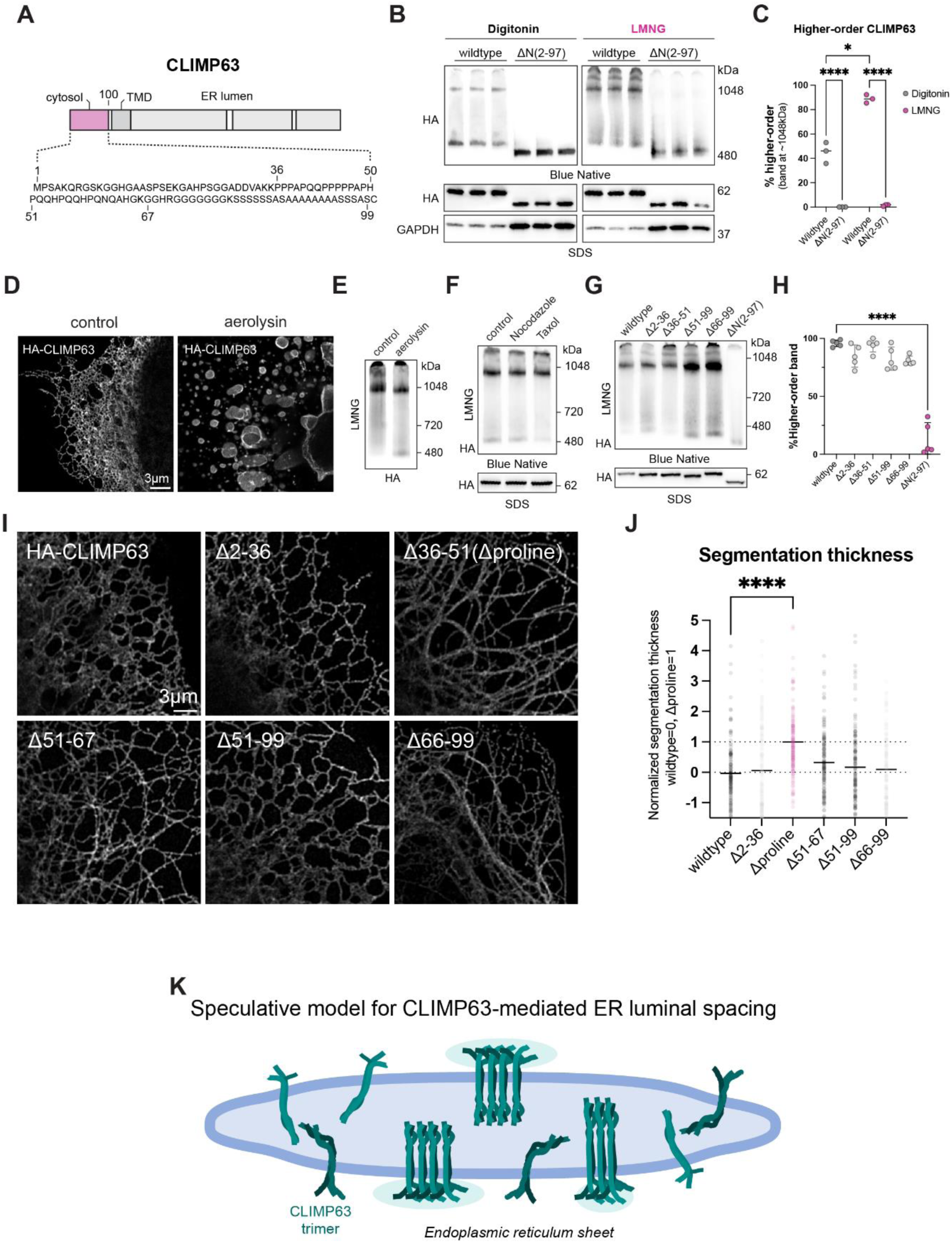
CLIMP63 trimers associate into higher-order structures through their cytosolic tails. **A.** Cytosolic domain organization of CLIMP63. **B.** Western blot analysis of CLIMP63 migrated on Blue Native or SDS-PAGE gels expressed in shCLIMP63 HeLa cells transfected with HA-CLIMP63 or HA-CLIMP63 lacking the cytoplasmic tail (residues 0-97) after solubulization in Digitonin or Lauryl Maltose Neopentyl Glycol (LMNG). GAPDH was used as a loading control. **C.** Quantification of percentage higher-order band (of total signal) at ∼1048kDa. Error bars represent mean ± standard deviation (n=3). *: p<0.01, ****: p<0.0001. **D.** Immunofluorescence (Aeryscan) of shCLIMP63 HeLa cells expressing human HA-CLIMP63 treated with 100ng/ml pro-aerolysin for 45 minutes, immunolabelled for HA. Scale bar: 3 μm. **E.** Western blot analysis of **(d)** migrated on Blue Native PAGE. **F.** Western blot analysis of HA-CLIMP63 migrated on Blue Native or SDS-PAGE gels expressed in shCLIMP63 HeLa cells transfected with HA-CLIMP63 and treated with 10μg/ml Taxol or Nocodazole for 2 hours. Control lane was treated with equivalent volume of DMSO. **G**. Western blot analysis of HA-CLIMP63 truncation mutants migrated on Blue Native or SDS-PAGE gels expressed in shCLIMP63 HeLa cells. **H**. Quantification of percentage higher-order band (of total signal) at ∼1048kDa. **I.** Immunofluorescence. (Aeryscan) of shCLIMP63 HeLa cells expressing HA-CLIMP63 and truncation mutatons immunolabelled for HA. Error bars represent mean ± standard deviation (n=5). ****: p<0.0001 **J.** Quantification of ER thickness of each mutant. Normalized thickness widltype=0, wildtype=0, Δproline=1 from **(I)**. Each dot represents the mean ER width of the image, obtained by segmenting the ER and averaging the ER width across all cells in that image. Error bars represent mean ± standard deviation (n=144 images per condition). ****: p<0.0001

Blue Native PAGE classically relies on digitonin, a mild non-ionic steroidal glycoside detergent that preserves many native protein-protein interactions during solubilization. We compared digitonin with LMNG (lauryl maltose neopentyl glycol), a synthetic non-ionic detergent of the neopentyl glycol class bearing two maltose head groups and two lauryl chains, which has an extremely low critical micelle concentration and is considered exceptionally mild for preserving protein complex integrity. Solubilization with LMNG led to a striking increase in the abundance of the ∼1,048 kDa species relative to the ∼480 kDa species (Figure 3B and C). The sensitivity of the higher-order assembly to detergent conditions indicates that the interactions mediating it are relatively weak and labile.

We next investigated the structural basis of this higher-order assembly. To test whether trans interactions of CLIMP63 trimers across the ER lumen contribute, we treated cells with the pore-forming toxin aerolysin. Aerolysin inserts into the plasma membrane and forms pores through which ions flow into and out of the cell, triggering vacuolation of the ER and expansion of the luminal space [26]. If higher-order CLIMP63 assembly depended on trans contacts across the lumen, this expansion would be expected to disrupt it. However, CLIMP63 migration on Blue Native PAGE was unchanged in aerolysin-treated cells relative to untreated controls (Figure 3D and 3E), arguing against trans luminal interactions as a driver of higher-order assembly. Since CLIMP63^LD^, which lacks the transmembrane and cytoplasmic domains, migrated as a single ∼480 kDa species (Figure S1), the higher-order assembly requires elements outside the luminal domain. To determine whether the cytoplasmic tail is responsible, we generated a deletion mutant lacking residues 1–97 (CLIMP63^ΔN^), which removes the cytoplasmic tail while retaining the transmembrane domain. CLIMP63^ΔN^ migrated as a single ∼480 kDa species irrespective of whether digitonin or LMNG was used (Figure 3B and 3C), demonstrating that the cytoplasmic tail is required for higher-order assembly.

The cytoplasmic tail of CLIMP63 has multiple known functions: it binds microtubules, has been reported to interact with nucleic acids, and is subject to phosphorylation and S-acylation at cysteine 100. To determine which of these properties mediates higher-order assembly, we tested each systematically. Disruption of the microtubule network by taxol stabilization or nocodazole-induced depolymerization did not alter CLIMP63 migration on Blue Native gels, excluding microtubule binding as a determinant of assembly (Figure 3F). S-acylation was similarly dispensable: neither the C100A mutant nor a double-cysteine mutant promoting hyperacylation affected higher order assembly (Figure S2B) consistent with previsou findings [17], and the ΔN mutant retained S-acylation as confirmed by acyl-RAC assay, confirming that the cytoplasmic tail and S-acylation contribute independently to CLIMP63 behavior (Figure S2C).

Enzymatic digestion of nucleic acids using Benzonase, TurboNuclease, or RNase A also had no effect on CLIMP63 migration (Figure S2D), excluding nucleic acid binding as the basis of higher-order assembly.

We nextgenerated a series of partial truncation mutants. None altered the migration pattern on Blue Native PAGE (Figure 3G and H), yet several produced marked changes in ER architecture (Figure 3I and J). This dissociation between higher order assembly and morphology indicates that the cytoplasmic tail has functionally separable roles: the contribution to higher-order assembly appears to require the intact tail, while specific sub-regions, including those governing microtubule binding, make independent contributions to ER architecture that are not mediated through changes in oligomeric state.

**Figure S2.**
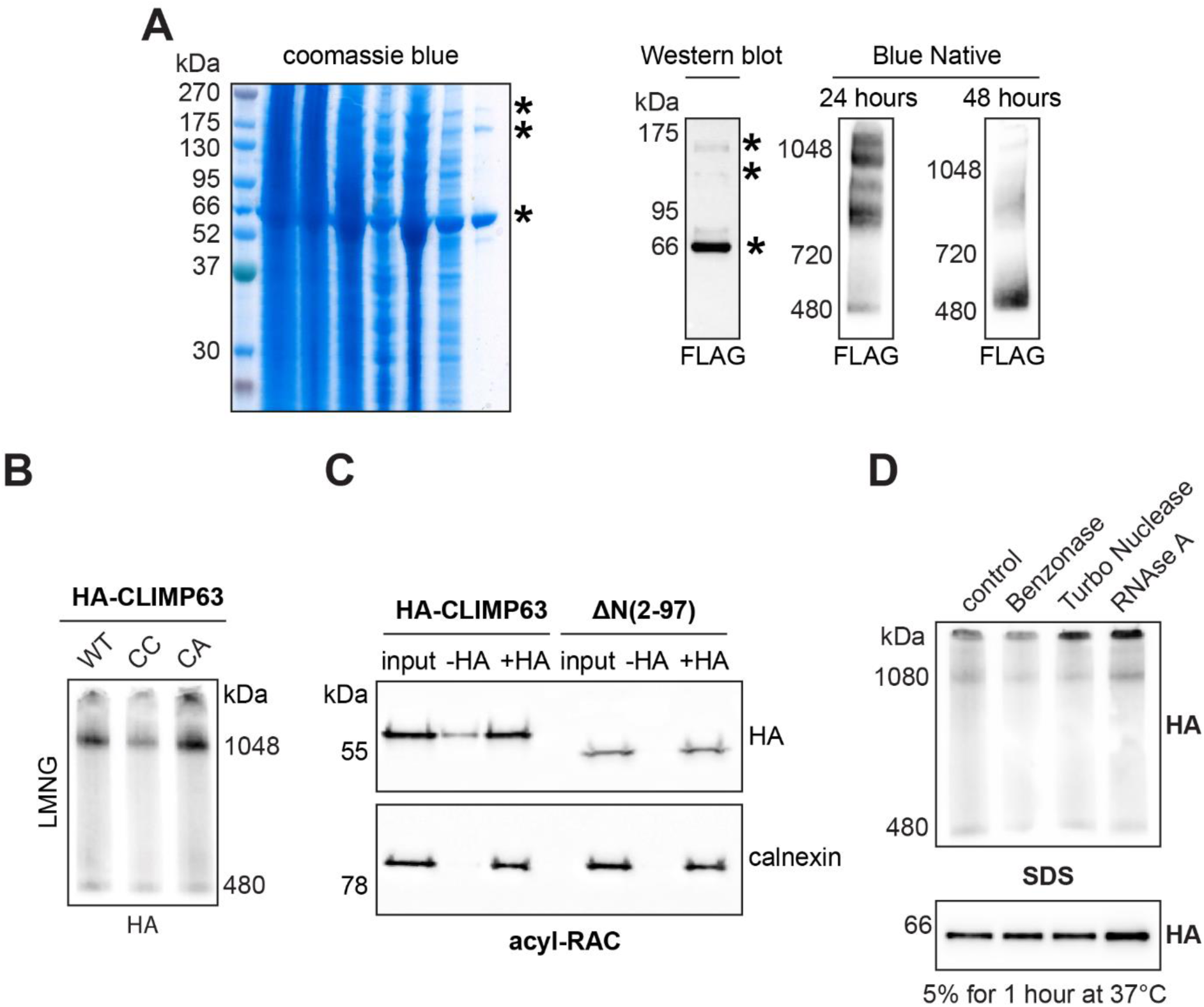
CLIMP63 trimers associate into higher-order structures through their cytosolic tails. **A.** Coomassie blue staining of purified His6x-FLAG-tagged full-length CLIMP63 solubulized in LMNG expressed in HEK293E cells. Western blot analysis of purified His6x-FLAG-tagged full-length CLIMP63. Western blot analysis of LMNG purified His6x-FLAG-tagged full-length CLIMP63 migrated on Blue Native 24 hours after purification and 48 hours after purification. **B.** Western blot analysis of HA-CLIMP63 and CC/CA mutants migrated on Blue Native gel expressed in shCLIMP63 HeLa cells. **C.** Acyl-rac assay of HA-CLIMP63 and HA-CLIMP63 lacking the cytosolic domain ΔN (residues 0-97) from shCLIMP63 HeLa cells following specific cleavage with hydroxylamine (+HA) (NH2OH). **D.** Western blot analysis of HA-CLIMP63 migrated on Blue Native or SDS-PAGE gels expressed in shCLIMP63 HeLa cells. Cells were treated with 5% DMSO as control, Benzonase, Turbo Nuclease, and RNAse A for 1 hour at 37°C, respectively, prior to LMNG solubulization.

## DISCUSSION

The data presented here revise the molecular model for CLIMP63-mediated ER shaping at several levels. CLIMP63 assembles into parallel, not antiparallel, trimers through its luminal coiled-coil domain. These trimers adopt an elongated rod-shaped conformation with regions of intrinsic conformational flexibility that are required for ER shaping. Higher-order assemblies beyond the trimer are driven by the cytoplasmic tail, and the cytoplasmic tail has functionally separable roles in oligomerization and ER architecture.

The antiparallel dimer model, in which CLIMP-63 molecules on opposing membranes interact tip-to-tip to hold the lumen at a defined width, was appealing for its simplicity. However, several observations reported here are inconsistent with it. The systematic cysteine-trapping data demonstrate a parallel orientation of CLIMP-63 chains within the trimer, which is incompatible with the antiparallel geometry. The absence of any effect of aerolysin-induced ER vacuolation on the Blue Native PAGE migration pattern argues against trans luminal interactions as a driver of the observed assemblies. The D528-STOP truncation, which removes residues proposed to mediate trans contacts, does not alter higher-order oligomerization. And the secreted luminal domain alone does not form the ∼1,048 kDa species as evidenced by Blue Native analysis of secreted double and tripple cysteine-trapping mutants Thus this higher order assembly is anchored at the membrane and not driven by luminal domain interactions. Together, these findings argue that CLIMP63 does not function as a fixed-length molecular ruler bridging opposing membranes.

We propose an alternative model in which CLIMP-63 trimers projecting from the membrane surface into the lumen act as a molecular brush that governs sheet width through physical rather than stoichiometric means. The concept of a brush controlling biological spacing is well established [27,28]. Neurofilaments are the canonical example: intrinsically disordered sidearms projecting radially from the neurofilament backbone create an entropic brush that repels neighboring filaments and determines axonal caliber [29,30]. The spacing set by the brush depends on sidearm length, charge, and density, parameters that can be regulated, rather than on any fixed structural interaction. The physical principle is that a dense layer of flexible polymers resists compression: as two brush-coated surfaces approach, the conformational entropy of the projections decreases and generates a repulsive pressure that sets an equilibrium spacing.

CLIMP-63 trimers implement a structural variant of this principle. Rather than disordered sidearms, they present semi-flexible coiled-coil rods, ordered secondary structures interrupted by hinge regions that introduce conformational freedom. The brush is therefore not entropic in the classical polymer sense, but the underlying mechanism is analogous: projections from the membrane resist compression of the luminal space, and their physical properties, length, flexibility, and density, determine the equilibrium spacing.

The structural parallel to VAP-A at membrane contact sites supports the plausibility of this mechanism for ER transmembrane proteins [31]. De la Mora et al. showed that VAP-A, which projects a flexible coiled-coil from the ER into the cytoplasm, forms a brush-like array at membrane contact sites whose density directly controls intermembrane distance, ranging from ∼15 nm when few VAP-A molecules are engaged to ∼30 nm in high-density membrane ribbons [31]. As in CLIMP-63, the conformational flexibility of the VAP-A coiled-coil, conferred by two disordered linker regions, is integral to this behavior: it allows each molecule to explore variable extensions and orientations, generating the repulsive pressure that separates the membranes. VAP-A thus provides a direct demonstration that a coiled-coil based brush anchored in the ER membrane can control intermembrane spacing in a density-dependent manner. The key difference is topological: VAP-A acts on the cytoplasmic face and bridges two organelle membranes at contact sites, whereas CLIMP-63 acts in the lumen of a single ER sheet, with contributions potentially from both leaflets.

The functional importance of CLIMP-63 luminal domain flexibility is demonstrated by the chimeric CLIMP-63^hum-fish^ experiment. Replacing the human luminal domain, which is semi-flexible due to its hinge regions, with the more rigid milkfish ortholog causes complete ER vacuolation. The only documented luminal binding partner of CLIMP-63 is calumenin, which acts as a chaperone indirectly affecting luminal ER width; no luminal interactions between CLIMP-63 and other sheet proteins have been reported [14]. The vacuolation phenotype therefore most directly implicates the mechanical properties of the coiled-coil itself. In the brush model, a more rigid rod would resist conformational compression more strongly, generating a repulsive pressure that expands rather than maintains the luminal space, a prediction consistent with the observed vacuolation. The hinge regions in the human CLIMP63 coiled-coil are thus not structural imperfections but functional elements that calibrate the brush to the physiological ∼50 nm set-point. This places CLIMP-63 in a growing class of proteins, alongside VAP-A, golgins, and the membrane contact site tether Ist2 [32–34], where conformational flexibility is an active determinant of membrane spacing rather than simply an absence of rigidity.

The lateral organization and density of the brush is likely governed by the cytoplasmic tail. The higher-order assembly of CLIMP63 trimers requires the intact cytoplasmic tail and is independent of microtubule binding, S-acylation, and nucleic acid interactions. This assembly is sensitive to the detergent used for solubilization, suggesting that the interactions are weak and labile, properties consistent with a dynamic, density-responsive array rather than a static lattice. Partial truncations of the tail that do not abolish higher-order assembly nevertheless alter ER morphology, indicating that the tail contributes to ER architecture through distinct, separable mechanisms. Clusters of VAP-A at membrane contact sites similarly arise from a combination of protein-protein interactions and geometric constraints rather than from a single deterministic mechanism [31], suggesting that dynamic lateral organization may be a general feature of brush-forming ER membrane proteins.

The brush model remains to be tested. Direct visualization of CLIMP-63 organization within native ER sheets by in situ cryo-electron tomography will be needed to determine whether trimers from opposing membranes are randomly distributed or adopt preferred relative orientations, and to establish whether the luminal projections interpenetrate or remain separated. Resolving these questions will define whether the CLIMP63 brush operates as two independent layers or as a single integrated structure, and will ultimately determine how molecular properties at the nanometer scale are transduced into the defined luminal geometry of the ER sheet.

## Methods

### Cell culture, transfections and drug treatments

All cells were grown in complete Dulbecco’s MEM, GlutaMAX (DMEM, Sigma) at 37 °C, supplemented with 10% foetal bovine serum (FBS), penicillin and streptomycin. Cells were mycoplasma negative as tested on a trimestral basis using the MycoProbe Mycoplasma Detection Kit (CUL001B). For transfection, cells were dissociated using trypsin and plated in tissue culture dishes (Falcon, US). After 24 h, the medium was changed and the cells were transfected using TransIT reagent (Mirus Bio). The cells were incubated for 24–48 h before performing experiments. Drug treatments were used at: pro-aerolysin (100 ng/mL), nocodazole (2 h at 10 µg/mL), taxol (2 h at 10 µg/mL) in complete medium.

### Protein production and purification

Suspension-adapted HEK293E cells were transiently transfected with constructs expressing the CLIMP63 luminal domain with an N-terminal signal secretion peptide and either an N- or C-terminal His6×-FLAG tag, using PEI MAX (Polysciences) in RPMI-1640 (Gibco) supplemented with 0.1% Pluronic F-68. After 1.5 h, cells were diluted into Excell293 medium (Sigma) supplemented with 4 mM glutamine and 3.75 mM valproic acid and agitated at 37 °C. Following a 7-day incubation, the cell culture medium was harvested by centrifugation and clarified using a 0.22 µm filter. The conditioned medium was purified by Ni-NTA affinity chromatography via the His-tag, or by anti-FLAG M2 affinity gel, followed by gel-filtration chromatography in 250 mM NaCl, 25 mM HEPES pH 7.5.

Full-length CLIMP63 bearing either an N- or C-terminal His6×-FLAG tag was expressed in suspension-adapted HEK293E cells. Cells were harvested by centrifugation at 3,000 × g for 30 min at 4 °C, washed in 1× PBS, and pelleted again. The pellet was resuspended in solubilization buffer (250 mM NaCl, 50 mM HEPES pH 7.5, 1% LMNG) supplemented with cOmplete™ EDTA-free Protease Inhibitor Cocktail (Roche; 1 tablet per 10 mL), homogenized manually in a Dounce homogenizer, and left to rotate at 4 °C for 2 h to extract membrane proteins. The lysate was clarified by ultracentrifugation at 4 °C. The supernatant, containing soluble His6×-FLAG-tagged CLIMP63, was incubated with anti-FLAG M2 affinity gel (Millipore, Billerica, MA) pre-equilibrated in wash buffer (250 mM NaCl, 25 mM HEPES pH 7.5). The beads were washed with 100 mL of wash buffer and eluted in the same buffer supplemented with 3×FLAG peptide. The eluate was concentrated using Amicon filters (Millipore, Billerica, MA) for cryo-EM grid preparation or for western blot analysis by Blue Native PAGE or SDS-PAGE.

### Immunofluorescence

Cells were seeded on glass coverslips (N1.5, Marienfeld, D) for at least 24 h. Fixation and permeabilization were optimised to preserve the cytoskeleton and ER membranes. Cells were washed 3× with PBS, followed by fixation using either (i) pre-cooled methanol for 4 min at −20 °C and washed 3× with PBS, or (ii) 3% PFA for 10 min at room temperature followed by saponin solubilization. The cells were then blocked overnight at 4 °C (or 30–60 min at RT) in PBS + 0.5% BSA (GE Healthcare, US). The coverslips were incubated with primary antibody for 30–120 min at RT, washed 3× for 5 min with PBS + 0.5% BSA, incubated for 30–45 min at RT with secondary fluorescent antibodies (Alexa 488, 568 or 647, Invitrogen, US), and finally washed again 3× with PBS + 0.5% BSA prior to mounting in ProLong mounting medium (ThermoFisher Cat#P36934). Coverslips were imaged by a LSM980 Airyscan microscope (Zeiss) with a 63× oil immersion objective (NA 1.4). Images were acquired using ZEN 2009 or ZEN Blue.

### Immunoprecipitation and Western blotting

For immunoprecipitation and western blot, cells were lysed on ice for 30 min with lysis buffer (50 mM Tris–HCl pH 7.4, 2 mM benzamidine, 10 mM NaF, 20 mM EDTA, 0.5% NP-40, and a protease inhibitor cocktail (Roche, CH)). The lysate was clarified by centrifugation at 4 °C for 3 min at 5,000 rpm. Lysates were pre-cleared using Sepharose G-beads alone for 30 min at 4 °C before immunoprecipitation (G-beads plus antibody) on a wheel overnight at 4 °C. The beads were then washed 3× with lysis buffer before adding 4× sample buffer containing β-mercaptoethanol. The samples were boiled for 5 min at 95 °C and vortexed before loading and migration on 4–12% or 4–20% Tris-glycine SDS-PAGE gels. Blots were revealed using a Fusion Absolute (Vilber Lourmat, CH) and quantified with ImageJ.

### Blue Native PAGE

Cells from 6 cm plates were washed 3× with PBS 24 h post-transfection and placed on ice. Cells were scraped, resuspended in 1 mL PBS, and centrifuged for 10 min at 10,000 × g at 4 °C. The pellet was dissolved in 100 µL cold lysis buffer (1% digitonin or 1% LMNG, NativePAGE Sample Buffer, and protease inhibitor cocktail (Roche, CH)). The lysate was passed through a 26 G needle and incubated on ice for 10 min on a rotating wheel, then clarified by centrifugation for 30 min at 20,000 × g at 4 °C. The supernatant was transferred and supplemented with G-250 at one-quarter the detergent concentration. After electrophoresis, gels were washed in 0.1% SDS for 15 min before dry western blot transfer.

### Acyl-RAC

Cells were lysed in Buffer A (0.2% SDS, 0.5% Triton X-100, 100 mM HEPES, 1 mM EDTA, and a protease inhibitor cocktail (Roche, CH)). Proteins were reduced with 25 mM TCEP for 1 h at RT and then blocked with 50 mM for 2 h at 40 °C. Proteins were precipitated with cold acetone at −20 °C for 20 min and centrifuged at 4 °C for 10 min at 7,500 rpm. The pellet was washed 5× with 70% acetone. After drying, samples were resuspended in SDS buffer. 10% of the sample was reserved as input and the rest was split into two tubes. The first tube was treated with 0.5 M hydroxylamine (final, in Tris pH 7.4) and 10% thiopropyl sepharose beads (Sigma). The second tube (negative control) received only Tris-HCl pH 7.4 with 10% thiopropyl sepharose beads. The samples were incubated at RT overnight. Finally, the beads were washed 3× in SDS buffer before adding 4× sample buffer with β-mercaptoethanol and performing SDS-PAGE followed by western blot as described above.

### Surface biotinylation

Cells were cooled with shaking at 4 °C for 15 min to arrest endocytosis. Cells were then washed three times with cold PBS and treated with EZ-Link Sulfo-NHS-SS-Biotin for 30 min with shaking at 4 °C. Cells were washed three times for 5 min with 100 mM NH4Cl and lysed in IP buffer for 15 min at 4 °C. Lysates were centrifuged for 5 min at 5,000 rpm and the supernatant incubated with streptavidin agarose beads overnight on a wheel at 4 °C. Beads were washed five times with IP buffer and proteins were eluted by incubation in SDS sample buffer with β-mercaptoethanol for 5 min at 95 °C, prior to SDS-PAGE and western blotting.

### Statistics and reproducibility

Statistical analyses were carried out using GraphPad Prism software. Data representation and statistical details can be found in the figure legends. Unless otherwise indicated, an unpaired two-tailed Student’s t-test was used for direct comparison of means between two groups, whereas ANOVA was used to compare means among three or more groups. For ANOVA analyses, p values were obtained by post hoc tests used to compare every mean or pair of means (Tukey’s and Sidak’s) or to compare every mean to a control sample (Dunnett’s). Data are represented as means ± standard deviation. ns: not significant, *p < 0.05, **p < 0.01, ***p < 0.001, ****p < 0.0001. All representative experimental data (e.g. western blots, Blue Native, immunofluorescence, and electron microscopy analysis) were repeated independently with equivalent results for a minimum of three biologically independent experiments.

